# Protocol for Simulating the Effect of THz Electromagnetic Field on Ion Channels

**DOI:** 10.1101/2024.09.09.612012

**Authors:** Lingfeng Xue, Zigang Song, Qi Ouyang, Chen Song

**Affiliations:** Center for Quantitative Biology, Academy for Advanced Interdisciplinary Studies, Peking University, Beijing 100871, China; School of Life Sciences, Peking University, Beijing 100871, China; School of Physics, Peking University, Beijing 100871, China; Peking-Tsinghua Center for Life Sciences, Academy for Advanced Interdisciplinary Studies, Peking University, Beijing 100871, China

## Abstract

Terahertz (THz) electromagnetic fields are increasingly recognized for their crucial roles in various aspects of medical research and treatment. Recent computational studies have demonstrated that THz waves can modulate ion channel function by interacting with either the channel proteins or the bound ions through distinct mechanisms. Here we outline a universal simulation protocol to identify the THz frequencies that may affect ion channels, which consists of frequency spectrum analysis and ion conductance analysis. Following this protocol, we studied the effect of THz field on a Ca_V_ channel and found a broad frequency band in 1 to 20 THz range. We believe that this protocol, along with the identified characteristic frequencies, will provide a theoretical foundation for future terahertz experimental studies.

## Main

Terahertz (THz, 10^12^ Hz) electromagnetic field is an emerging technology with significant potential in medical diagnostics and treatment. ^1^ As an imaging modality, THz waves have been employed to detect various types of cancers, including breast tumors and skin cancer.^2,3^ Additionally, THz technology has been explored for the diagnosis and treatment of a range of diseases such as thyroid nodules, Alzheimer’s disease, etc. ^4^

The biological effects of THz fields can be broadly categorized into thermal and non-thermal effects. The thermal effect arises from the absorption of THz radiation by biological tissues, resulting in the conversion of electromagnetic energy into heat. In contrast, the non-thermal effects encompass a range of biological responses. For instance, *in vitro* studies have shown that THz radiation can induce morphological abnormalities in neuron cell membranes and alter intracellular structures. ^5^ Additionally, when THz radiation was applied to mammalian stem cells, certain genes were activated while others were repressed, suggesting its potential role in cellular reprogramming. ^6^ Mathematical modeling studies have also indicated that THz radiation may induce resonance effects in DNA, leading to modifications in gene expression. ^7^ These effects are thought to be mediated by the fact that many biomolecules exhibit inter- and intra-molecular motions within the THz frequency range, contributing to both the thermal and non-thermal interactions of THz fields.

Recently, several groups have investigated the effects of THz fields on ion channels. For example, it was found that a 5.6 ½m (53.5 THz) field can increase potassium current ^8^(Figure 1A). Molecular dynamics (MD) simulations on the KcsA channel revealed that a THz field with a frequency of 51.87 THz significantly increased potassium current, which corresponds to the vibration of -C=O groups in the selectivity filter (SF) region. ^9^ Similarly, MD simulations on the Ca_V_Ab channel showed that a frequency of 42.55 THz enhanced the channel’s permeability, corresponding to the stretching mode of the carboxylate group at the SF.^10^ These findings indicate that THz fields can modulate ion channels by affecting specific functional groups of proteins (Figure 1B).

**Figure 1:**
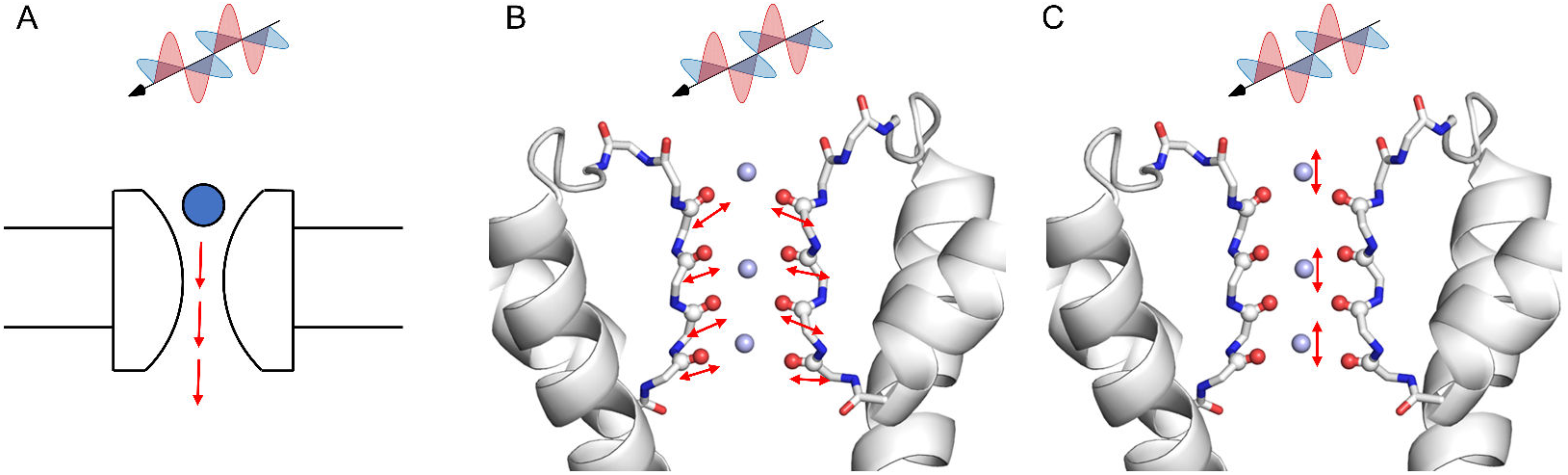
Effect of THz fields on ion channels through different mechanisms. (A) THz fields can affect the ion conductance. (B) THz fields affect the vibration motions of carbonyl bonds at the selectivity filter (SF). (C) THz fields affect the vibration motions of ions at the SF.

Our recent studies further demonstrated that the ion oscillation frequencies also lie within the THz range, and the application of specific THz frequencies can regulate ion conductance. ^11^ Based on these findings, we proposed a new mechanism for the regulation of ion channels (Figure 1C), wherein the applied THz field directly influences ion motion, accelerating the ions as they permeate the channel. It was unclear whether the enhanced ion conductance was due to a thermal effect. To investigate this, we applied electric fields in different directions. We found that ion conductance increased only when the THz field was applied in the z-direction (perpendicular to the membrane), whereas fields applied in the x- or y-directions had no significant effect. This direction-dependence indicates that the observed enhancement of ion conductance by THz fields is not simply a thermal effect, thereby further sparking interest in the study of THz interactions with ion channels.

To advance the study of THz effects on ion channels, here we summarize a universal simulation protocol designed to identify THz frequencies that may influence ion conductance (Figure 2). This protocol includes the following steps:

**Figure 2:**
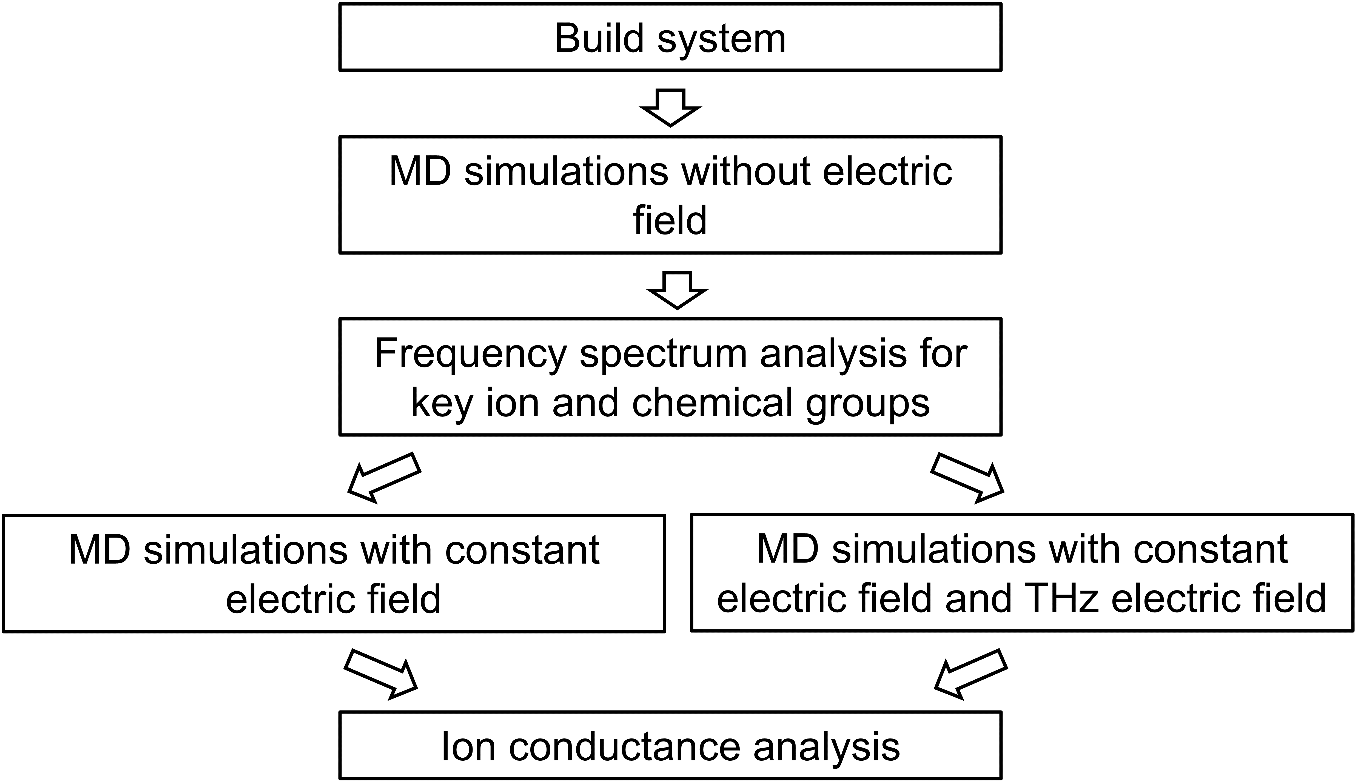
The overall simulation and analysis protocol for studying the effect of THz fields on ion channels.

1. Select the ion channel of interest, construct the MD simulation system, and perform adequate equilibrium simulations. Charmm-GUI is a convenient tool for this step. ^12^ An open-state channel is preferred, as we may want to validate the effect of the THz field by simulating the conductance of the channel in the fourth step.
2. Initiate MD simulations without any applied electric field. During these simulations, the velocities of all the atoms, or the atoms of interest, are sampled at an interval of 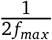, where *f*_*max*_ is the highest frequency in the intended spectrum. In our case, the highest frequency is 100 THz, so the velocities are recorded every 5 fs, allowing the high-frequency motions to be fully captured.
3. Perform frequency spectrum analysis on the velocities of key ions and protein functional groups to identify their characteristic frequencies. This can be accomplished using FFT (Fast Fourier Transform) analysis. For instance, one can use the numpy.fft function in Python to execute the analysis. It is crucial to pay special attention to the ion-binding sites along the permeation pathway. This includes not only the chemical groups of the ion channels that form these binding sites but also the bound ions themselves, as these often exhibit motions that fall within the THz frequency range, making them likely targets for modulation by THz fields.
4. Using these characteristic frequencies, we applied THz fields to assess their impact on ion conductance. This step can be further divided into two parts:
  - Simulate the ion conduction in the absence of the THz field. We apply a constant electric field in the z-direction (membrane normal) to calculate the conductance of the channel, using the formula 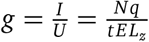, where *g* is conductance, *I* is current, which is calculated by the number of ion permeation events *N* for a given period *t, U* is transmembrane voltage, which is proportional to the electric field *E* and the z-direction box size *L*_*z*_. The calculated conductance should be compared to the experimental values to validate the simulation parameters and protocol.
  - Simulate ion conduction in the presence of the THz field by applying an additional THz electric field, in addition to the constant electric field in the z-direction. The magnetic field can be neglected due to its minimal impact compared to the electric field on ion channels. Apply the THz field in the x, y, and z directions, respectively. As the current GROMACS package does not fully support an additional THz field along the same direction of the constant electric field, we developed a plugin to introduce an extra constant electric field (https://github.com/ComputBiophys/GromacsPlugin). Using this plugin, perform simulations with both THz and constant electric fields to determine the ion conductance of the channel.
  - Compare the simulated ion conductances from the simulations with and without the THz field to assess whether the selected THz frequency can significantly alter ion conductance. It is important to note that both the frequency and the direction of the THz field are crucial factors, likely corresponding to the motion characteristics of the chemical groups or ions analyzed in the third step above.

Overall, this computational protocol allows us to identify the characteristic frequencies of key ions and functional groups, as well as to evaluate the effects of THz fields on channel conductance at these frequencies.

Using the above protocol, we calculated the THz frequencies for the Ca_V_1.3 channel (Figure 3). We utilized a closed-state Ca_V_1.3 structure ^13^ for the simulations, as an open-state structure is not currently available. To assess model dependency, we adopted two different calcium models: the default CHARMM model and our recently developed multi-site calcium model. ^14^ Following the protocol steps 1-–3, we identified the characteristic frequencies for Ca^2+^ ions at the SF to be in the range of 1 to 20 THz for both models (Figure 3A, C). With the CHARMM calcium model, the peak frequency was approximately 8 THz, while simulations using the multi-site calcium model revealed peak frequencies at 2, 5, and 16 THz. Although the results from the two models were not identical, both indicate that THz frequencies in the range of 1–20 THz may influence Ca^2+^ conductance. Note that the artificial peak at 69 THz, observed with the multi-site calcium model, was attributed to bond vibrations of pseudo atoms and should be disregarded. For the carboxylate groups in the SF of Ca_V_1.3, we identified two characteristic frequencies at 42.6 and 48.5 THz (Figure 3B, D), which are very close to previous results observed in Na_V_ channels. Ion permeation simulations (steps 4—5) were not conducted due to the unavailability of an open-state structure for Ca_V_.

**Figure 3:**
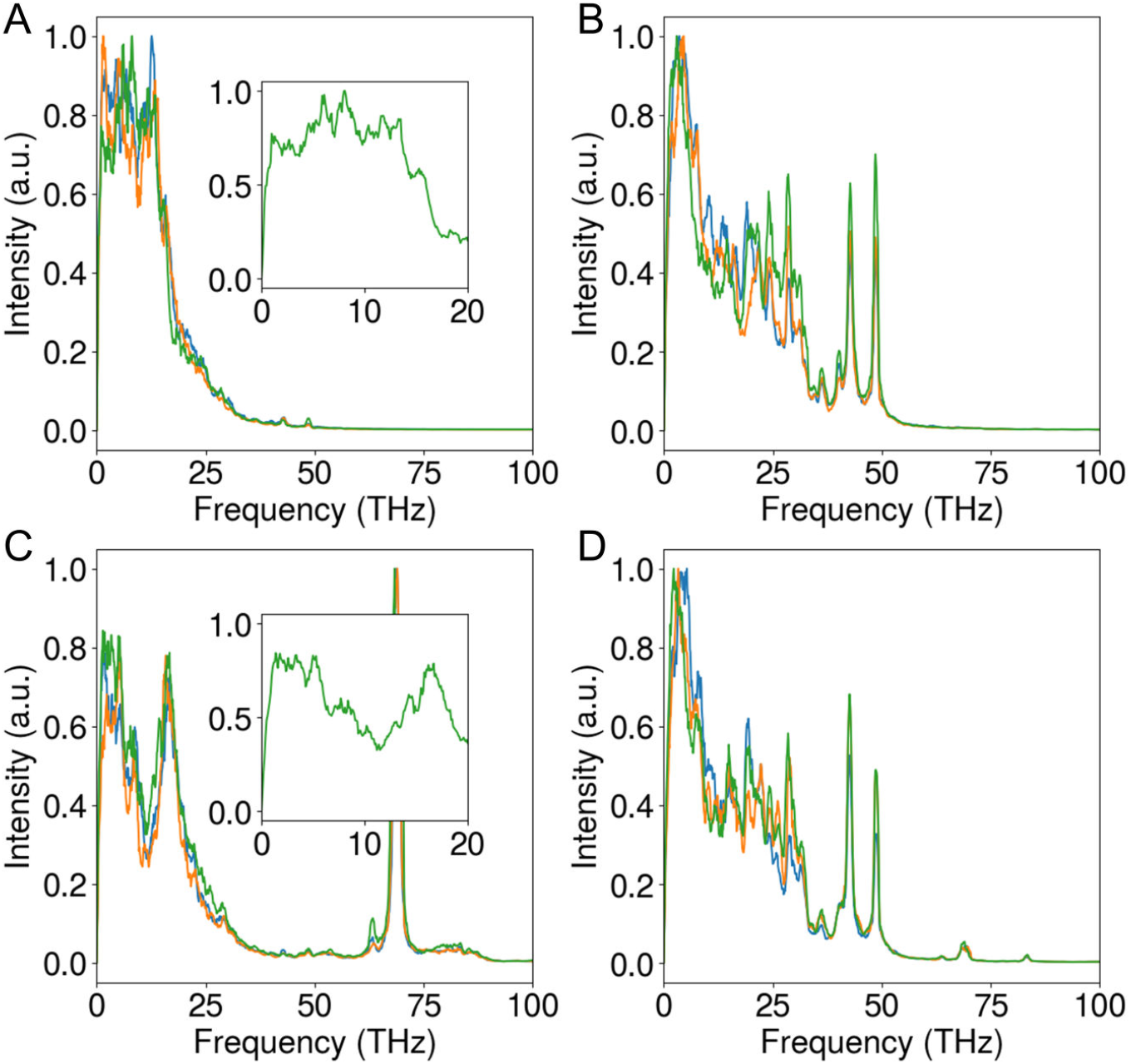
Frequency spectrum of Ca^2+^ ions (A) and carboxylate oxygens (B) at the selectivity filter of Ca_V_1.3 with default CHARMM calcium model. (C,D) Similar to A–B, but obtained with the multisite calcium model. The blue, orange, and green lines represent the frequency spectra for velocities in the x, y, and z directions respectively.

In summary, we present a table (Table 1) detailing the likely characteristic THz frequencies and the corresponding chemical groups of voltage-gated ion channels that may be affected. For K^+^ ions in K_V_1.2, the characteristic frequencies are 1.4, 2.2, and 2.9 THz. For Na^+^ ions in Na_V_1.5, the characteristic frequency range is broad, spanning 1–10 THz, with notable peaks at 2.5 and 10 THz. Similarly, for Ca^2+^ ions in Ca_V_1.3, the characteristic frequency ranges from 1–20 THz, with peaks at 2, 5, 8, and 16 THz. In addition, the chemical groups that bind the permeating ions in the SF of ion channels show important frequencies too, including the carbonyl bond stretching in K_V_1.2 with a frequency of 51 THz and the carboxylate bond stretching in sodium and calcium channels with frequencies of 42.5 and 48.6 THz.

**Table 1:** Characteristic frequencies of ions and chemical groups at the selectivity filter of ion channels.

| Channel | Characteristic frequency (THz) | Vibrational mode |
| --- | --- | --- |
| KcsA<sup>9</sup> | 51.87 | C=O bond stretching |
| Ca<sub>v</sub>Ab<sup>10</sup> | 42.55 | C-O bond stretching |
| K<sub>v</sub>1.2<sup>11</sup> | 1.4, 2.2, 2.9 | K<sup>+</sup> vibration in z direction |
|  | 5.0 | K<sup>+</sup> vibration in x/y direction |
|  | 10.8 | C=O wiggling in z direction |
|  | 51 | C=O bond stretching |
| Na<sub>v</sub>1.5<sup>11</sup> | 2.5, 10 | Na<sup>+</sup> vibration |
|  | 42.5, 48.6 | C-O bond stretching |
| Ca<sub>v</sub>1.3 | 2, 5, 8, 16 | Ca<sup>2+</sup> vibration |
|  | 42.6, 48.5 | C-O bond stretching |

Despite the insights gained, computer simulations have limitations. For example, the electric field strengths used in simulations are considerably higher than those in experimental conditions, and results may vary depending on the force field employed. Nonetheless, the calculated frequencies offer valuable references for future experimental studies and could contribute to advancing THz biology.

## Acknowledgements

The MD simulations were performed on the computing platform of the Center for Life Sciences at Peking University and the National Supercomputer Center in Tianjin.

## Code and data availability

The MD plugin can be found on GitHub (https://github.com/ComputBiophys/GromacsPlugin). The simulation data are available upon reasonable request.

## References

[1] Amini, T.; Jahangiri, F.; Ameri, Z.; Hemmatian, M. A. A review of feasible applications of THz waves in medical diagnostics and treatments. Journal of Lasers in Medical Sciences 2021, 12, e92.

[2] Fitzgerald, A. J.; Wallace, V. P.; Jimenez-Linan, M.; Bobrow, L.; Pye, R. J.; Purushotham, A. D.; Arnone, D. D. Terahertz pulsed imaging of human breast tumors. Radiology 2006, 239, 533–540.

[3] Woodward, R. M.; Cole, B. E.; Wallace, V. P.; Pye, R. J.; Arnone, D. D.; Linfield, E. H.; Pepper, M. Terahertz pulse imaging in reflection geometry of human skin cancer and skin tissue. Physics in Medicine & Biology 2002, 47, 3853.

[4] Shi, S.; Yuan, S.; Zhou, J.; Jiang, P. Terahertz technology and its applications in head and neck diseases. IScience 2023, 26.

[5] Olshevskaya, J.; Ratushnyak, A.; Petrov, A.; Kozlov, A.; Zapara, T. Effect of terahertz electromagnetic waves on neurons systems. 2008 IEEE Region 8 International Conference on Computational Technologies in Electrical and Electronics Engineering. 2008; pp 210–211.

[6] Bock, J.; Fukuyo, Y.; Kang, S.; Phipps, M. L.; Alexandrov, L. B.; Rasmussen, K. Ø.; Bishop, A. R.; Rosen, E. D.; Martinez, J. S.; Chen, H.-T.; others Mammalian stem cells reprogramming in response to terahertz radiation. PloS one 2010, 5, e15806.

[7] Alexandrov, B. S.; Gelev, V.; Bishop, A. R.; Usheva, A.; Rasmussen, K. Ø. DNA breathing dynamics in the presence of a terahertz field. Physics Letters A 2010, 374, 1214–1217.

[8] Liu, X.; Qiao, Z.; Chai, Y.; Zhu, Z.; Wu, K.; Ji, W.; Li, D.; Xiao, Y.; Mao, L.; Chang, C.; others Nonthermal and reversible control of neuronal signaling and behavior by midinfrared stimulation. Proceedings of the National Academy of Sciences 2021, 118, e2015685118.

[9] Wang, Y.; Wang, H.; Ding, W.; Zhao, X.; Li, Y.; Liu, C. Regulation of ion permeation of the KcsA channel by applied midinfrared field. International Journal of Molecular Sciences 2022, 24, 556.

[10] Li, Y.; Chang, C.; Zhu, Z.; Sun, L.; Fan, C. Terahertz wave enhances permeability of the voltagegated calcium channel. Journal of the American Chemical Society 2021, 143, 4311–4318.

[11] Song, Z.; Xue, L.; Ouyang, Q.; Song, C. The Effect of THz Electromagnetic Field on the Conductance of Potassium and Sodium Channels. bioRxiv 2024,

[12] Jo, S.; Kim, T.; Iyer, V. G.; Im, W. CHARMM-GUI: a web-based graphical user interface for CHARMM. Journal of computational chemistry 2008, 29, 1859–1865.

[13] Yao, X.; Gao, S.; Yan, N. Structural basis for pore blockade of human voltage-gated calcium channel Cav1. 3 by motion sickness drug cinnarizine. Cell Research 2022, 32, 946–948.

[14] Zhang, A.; Yu, H.; Liu, C.; Song, C. The Ca2+ permeation mechanism of the ryanodine receptor revealed by a multi-site ion model. Nature Communications 2020, 11, 922.

